# Changes in natural transformation after salt adaptation

**DOI:** 10.1101/2021.12.05.471276

**Authors:** HA Kittredge, SE Evans

## Abstract

The exchange of genes between potentially unrelated bacteria is termed horizontal gene transfer (HGT) and is a driving force in bacterial evolution. Natural transformation is one mechanism of HGT where extracellular DNA (eDNA) from the environment is recombined into a host genome. The widespread conservation of transformation in bacterial lineages implies there is a fitness benefit. However, the nature of these benefits and the evolutionary origins of transformation are still unknown. Here, I examine how ∼330 generations or 100 days of serial passage in either constant or increasing salinities impacts the growth rate and transformation efficiency of *Pseudomonas stutzeri*. While the growth rate generally improved in response to serial transfer, the transformation efficiency of the evolved lineages varied extensively, with only 39-64% of populations undergoing transformation at the end of adaptive evolution. In comparison, 100% of the ancestral populations were able to undergo natural transformation. I also found that evolving *P. stutzeri* with different cell lysates (or populations of dead cells) minimally affected the growth rate and transformation efficiency, especially in comparison to the pervasiveness with which transformation capacity was lost across the evolved populations. Taken together, I show that the efficiency of eDNA uptake changes over relatively rapid timescales, suggesting that transformation is an adaptive and selectable trait that could be lost in environments where it is not beneficial.

## Introduction

Natural transformation is a mechanism of horizontal gene transfer whereby bacteria acquire extracellular DNA (eDNA) from the environment and recombine it into their genomes. Transformation plays a key role in bacterial evolution (Ambur et al. 2016; Koonin 2016); however, the fitness benefits of transformation remain unknown, despite extensive study. While it is generally accepted that transformation can facilitate adaptation through genetic recombination, the consequences of this genetic exchange can be both beneficial and costly (Baltrus 2013). The theoretical benefits of transformation are similar to meiotic sex and include speeding up adaptation, combining beneficial genes into one genome, and separating beneficial mutations from deleterious loads (Kim and Orr 2005; Cooper 2007; Mell and Redfield 2014; Takeuchi et al. 2014; Koonin 2016). However, extracellular DNA from dead bacteria can also carry an increased mutational load or promote the spread of selfish genes (Redfield 1988; Redfield et al. 1997). Consequently, the fitness advantages of transformation and the environmental conditions in which they are conferred have been difficult to quantify experimentally (see Table 1).

**Table 1.**
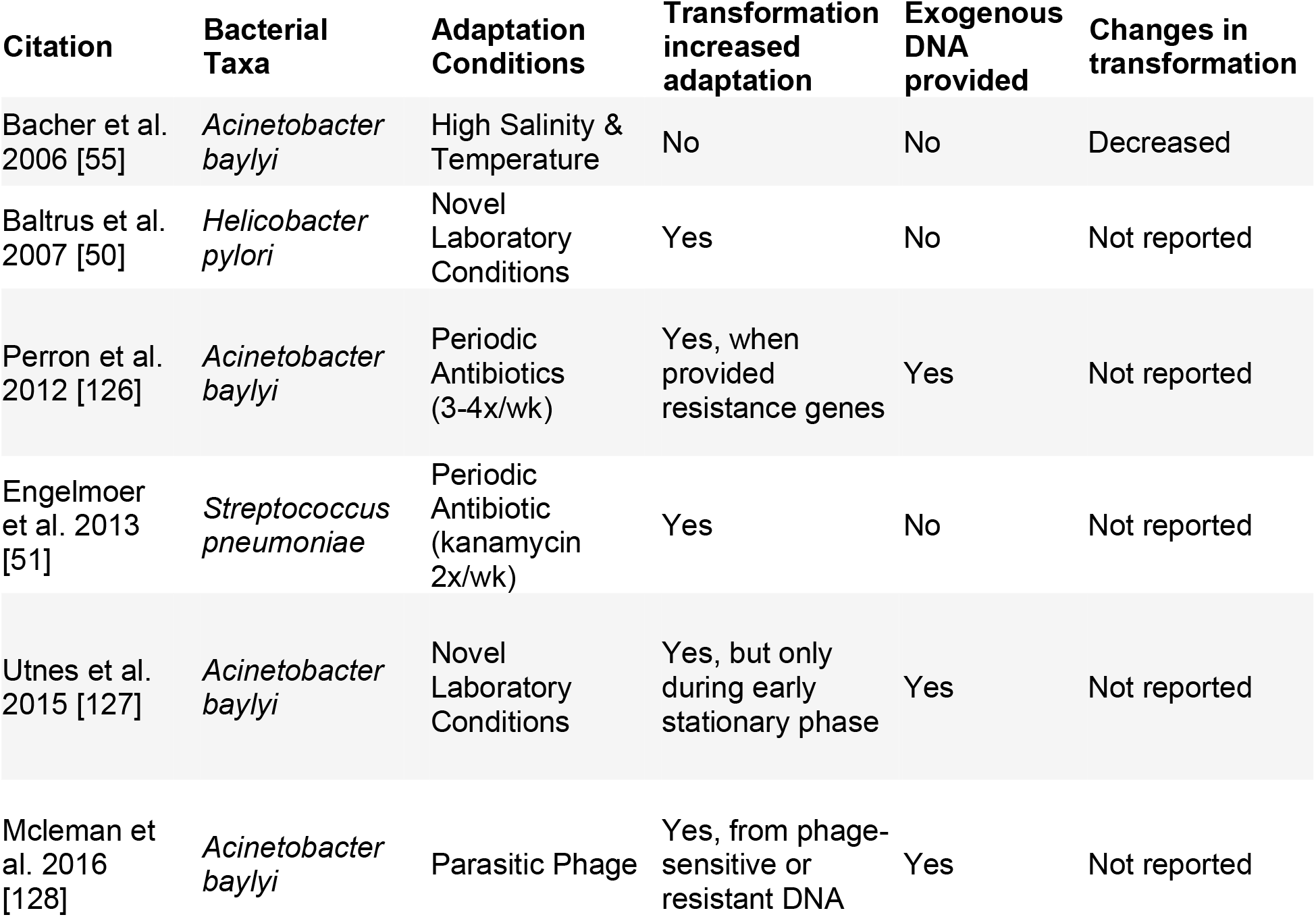
Review of transformation-mediated fitness effects. Experimental evolution studies that have quantified the fitness effects of transformation.

| Citation | Bacterial Taxa | Adaptation Conditions | Transformation increased adaptation | Exogenous DNA provided | Changes in transformation |
| --- | --- | --- | --- | --- | --- |
| Bacher et al. 2006 [55] | *Acinetobacter baylyi* | High Salinity & Temperature | No | No | Decreased |
| Baltrus et al. 2007 [50] | *Helicobacter pylori* | Novel Laboratory Conditions | Yes | No | Not reported |
| Perron et al. 2012 [126] | *Acinetobacter baylyi* | Periodic Antibiotics (3-4x/wk) | Yes, when provided resistance genes | Yes | Not reported |
| Engelmoer et al. 2013 [51] | *Streptococcus pneumoniae* | Periodic Antibiotic (kanamycin 2x/wk) | Yes | No | Not reported |
| Utnes et al. 2015 [127] | *Acinetobacter baylyi* | Novel Laboratory Conditions | Yes, but only during early stationary phase | Yes | Not reported |
| Mcleman et al. 2016 [128] | *Acinetobacter baylyi* | Parasitic Phage | Yes, from phage-sensitive or resistant DNA | Yes | Not reported |

There are several potential explanations for the evolutionary maintenance of transformation [reviewed in 42]. Several of them posit that transformation evolved as a byproduct of acquiring DNA for nutrients (Redfield 1993; Claverys et al. 2006; Moradigaravand and Engelsta 2013; Sinha et al. 2013) or genome repair (Michod et al. 1988; Charpentier et al. 2012; Johnston et al. 2014). However, the presence of cellular machinery dedicated to protecting extracellular DNA (eDNA) from degradation inside the cell, suggests that eDNA is not acquired purely for the nutrient benefit (Seitz and Blokesch 2013; Johnston et al. 2014). In addition, many bacterial taxa preferentially kill and transform eDNA from close relatives, a process somewhat analogous to the exchange of DNA in eukaryotic sex (Guiral et al. 2005; Veening and Blokesch 2017).

Transformation is also similar to eukaryotic sex in that it is primarily beneficial in stressful or continuously changing environments. Population genetic models (Lynch et al. 1993; Takeuchi et al. 2014; Koonin 2016) and experimental evolution studies (Baltrus et al. 2007; Engelmoer et al. 2013) have shown that transformation is beneficial in rapidly fluctuating or stochastic environments where transformable cells can outcompete non-transformers (Palmer and Cartwright 2018; Carvalho et al. 2019). Theoretically, this is because transformation can increase genetic variation, thereby increasing the efficiency of natural selection (Barton and Charlesworth 1998). Transformation does not always provide a fitness benefit in stressful environments though, as Bacher et al. (2006) found that competent lineages of *Acinetobacter baylyi* did not adapt to novel laboratory conditions faster than their non-competent competitors, and repeatedly lost the ability to transform eDNA.

While several other studies have shown that transformation is beneficial in stressful environments (see Table 1), it is still unclear how the availability of beneficial genes might alter transformation-mediated fitness effects. For instance, antibiotic resistance only evolved via transformation when antibiotic resistance genes were provided, while phage resistance evolved in the presence of phage-sensitive or -resistant DNA (Perron et al. 2012; Mcleman et al. 2016). Since sequence similarity improves the efficiency of homologous recombination, it is generally accepted that transformation is most prevalent between closely related organisms (Soucy et al. 2015). However, sharing genes with close relatives could limit the acquisition of novel gene combinations and ultimately limit adaptation.

Here, I aim to better understand the evolutionary benefits of transformation by evolving *Pseudomonas stutzeri* – a highly transformable soil bacterium – in either constant or increasing salt concentrations for 100 days, while supplying different sources of eDNA (cell lysates or dead cells). At the end of the experiment, I quantify the growth rate, population size, and transformation efficiency (transformants/μg eDNA) of the evolved populations and compare this to the same measurements in the starting isolate or ancestor. I specifically address the following questions: 1) Does the transformation efficiency increase in response to evolving in a variable – relative to a constant environment (increasing salinity vs. constant low salinity)? 2) Does evolving with dead halophiles or dead *Pseudomonad* relatives better facilitate adaptation to high salt concentrations?

## Methods

### Serial dilution experiment

We serially transferred *Pseudomonas stutzeri*, strain 28a24 for ∼330 generations (100 days) in 96-well microtiter plates [see 101 for whole genome sequence]. Cultures were serially transferred every 24hrs at a 1:10 dilution and maintained at 26°C. For the first 50 days (∼170 generations) of the experiment, all populations were transferred as one treatment in a constant salt media (1.5% salinity; 10g/L tryptone, 5g/L yeast extract, and 15g/L NaCl). After 50 days (∼170 generations), the experiment was shut down due to the global covid-19 pandemic, and populations preserved in 40% glycerol at 20°C. Four weeks later, populations were revived and serially passed at 1.5% salinity for 4 days before the experiment was ‘re-started’ on day 51 (∼170 generations). At this point, we split the experiment into two treatments. The original treatment was maintained at a low constant salinity for the remainder of the experiment (1.5% salt media from day 1 to 100). The new treatment, which we refer to as the increasing salinity treatment was transferred to a 2% salt media (20g/L NaCl), where it was serially passed for 100 generations. Then on day 81, we increased the salt concentration to 2.5%, were it stayed for 67 generations until Day 100 (see Fig. 1 for the serial transfer conditions). The constant low and increasing salinity treatments had 96 replicates each. In addition, during each transfer (every 24hrs), populations were supplemented with eDNA via whole populations of dead bacteria – which equated to 5ng of genomic eDNA each transfer. We refer to these as cell lysates as they contain DNA and other cellular components (see Table S1 for a detailed list of the cell lysate sources).

**Fig. 1.**
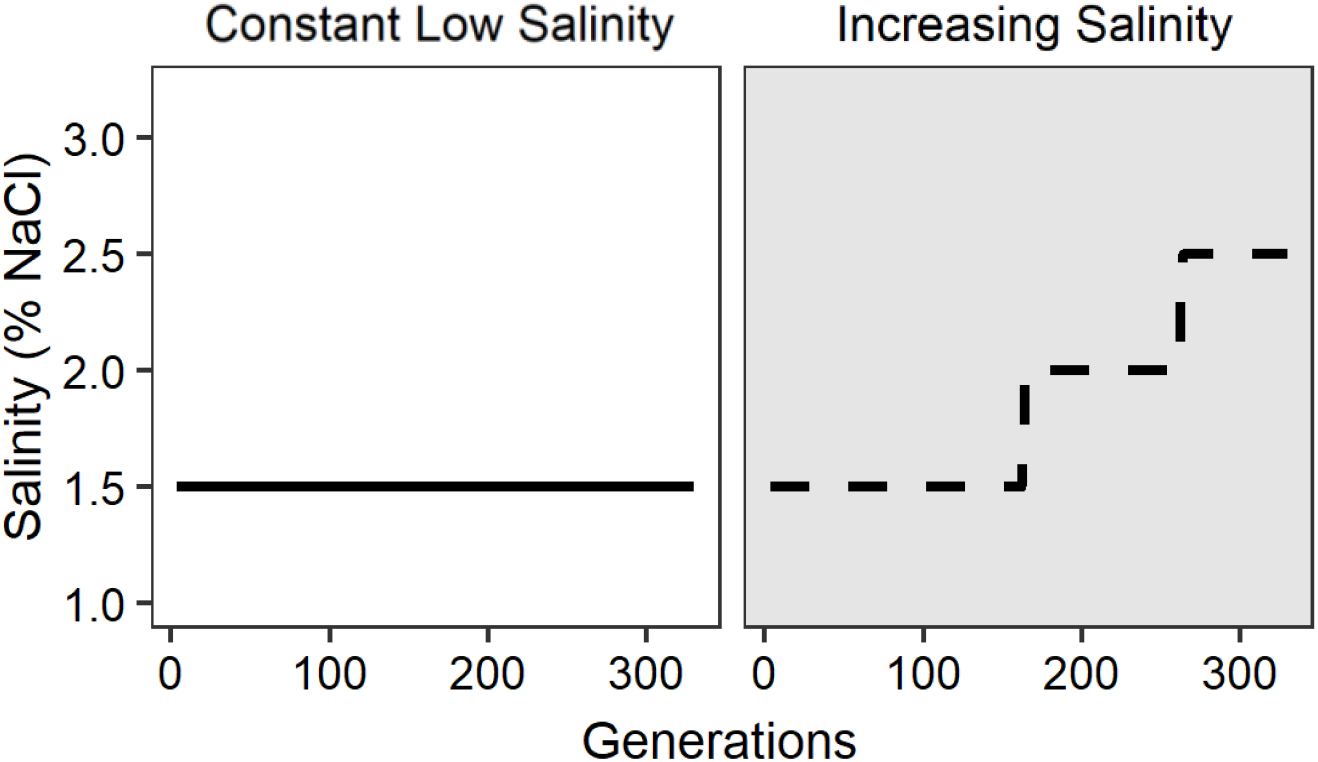
The serial transfer conditions for the two evolution treatments. The left panel shows the conditions for populations adapted to a constant low salinity environment (1.5% salinity) and the right panel shows the conditions for populations adapted to increasing salinities (1.5% to 2.5% salinity).

### Preparation of cell lysates

Individual cell lysates were prepared in 100mL batch cultures in liquid LB on a shaker table at 120rpm and 30°C (10g/L tryptone, 5g/L yeast extract, and 5g/L NaCl). After 48hrs, each culture was plated to confirm there was no contamination (10 μl replicate dots plated 3x). Each culture was then heat shocked at 90°C for 1hr. After heat shock, each culture was plated to confirm all the cells were dead. If bacterial strains still had viable colonies, these cultures went through another round of heat shock at 100-110°C, which was sufficient to kill the remaining cells. The heat shocked cultures were then spun down and resuspended in sterile nanopore water. The lysates were filtered through a 0.22μm filter and standardized to a concentration of 1ng DNA/μl using a Qubit 2.0 fluorometer (Life Technologies, USA). There were 12 cell lysate treatments with 8 replicates each. See Table S1 for expanded list of cell lysates.

### Growth rate determination

All assays were conducted on the ancestral population and populations that evolved for 100 days (∼330 generations). Strains of *P. stutzeri* were revived from 40% glycerol storage (−80°C) and diluted 1:10 in liquid LB media (0.5% salinity: 10g/L tryptone, 5g/L yeast extract, and 5g/L NaCl) in 250-μl microwell plates. After revival, the populations were transferred every 24hrs at a 1:10 dilution in 0.5% salinity for 4 transfers. After the fourth transfer, we moved the populations to two separate salt environments in 250-μl microwell plates, to quantify the growth rate and transformation capacity. In the constant low salinity treatment, 10 of the 96 populations never revived. In the increasing salinity treatment, 1 population never revived and 1 was contaminated, these are not reported in the results (see Table S2 for a full list).

Each 24hr assay was conducted at low salinity (1.5% salinity: 10g/L tryptone, 5g/L yeast extract, and 15g/L NaCl) and high salinity – a novel and stressful environment for *P. stutzeri* (3% salinity: 10g/L tryptone, 5g/L yeast extract, and 30g/L NaCl). For each population, we monitored absorbance at 600nm for 24hrs using a Biotek Synergy microplate reader (Winooski, VT). The growth curve data was fit to a standard form of the logistic equation using the Growthcurver package in R studio. We used the logistic equation to describe the population size N_t_ at time t:

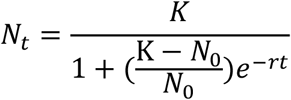

### Transformation efficiency and frequency

Cultures were revived following the same protocol used for growth rate determination. After revival, we quantified the transformation efficiency by tracking the acquisition of gentamicin resistance into the evolved and ancestral *P. stutzeri* populations which were gentamicin susceptible. The eDNA encoding gentamicin resistance was prepared from a mutant strain of *P. stutzeri*, strain 28a24, which carries a gentamycin resistance gene and LacZ gene fused to a miniTn7 transposon (Tn7 transposition of pUC18-mini-Tn7T-Gm-lacZ). To begin the assay, we transferred 20μl from each evolved and ancestral population into 180μl of fresh LB media containing 1.5% or 3% salinity. I added genomic extracellular DNA (eDNA) resuspended in nanopore water to each population and incubated at 30°C. After 24hrs we performed a serial dilution and titers were determined on selective media (LB + gentamycin [50 μg/ml] + Xgal [40 μg/ml]) and non-selective media (LB) using triplicate 10μl dots. Population level transformation efficiency was determined by dividing the average number of transformants in a population by the μg of eDNA (0.02μg). We also report population level transformation frequencies by dividing the average number of transformants by the total number of cells or the population size.

### Statistical analyses

Prior to analysis, we checked that data met assumptions of normality and homogeneity of variance. We corrected for increased homogeneity of variance across population sizes using a log transformation. We analyzed bacterial growth rates and populations sizes using two-factor ANOVA, with Evolution Conditions, Assay Conditions, and their interaction as factors. For each evolution treatment – constant salinity versus increasing salinity – we used a two-factor ANOVA with the Assay Conditions, Cell Lysate treatment, and their interaction as factors. we determined differences in transformation capacity using a general linearized model with a negative binomial distribution to account for positive skew. To determine statistical differences in the number of non-transforming populations (zeros) we used a two-part hurdle model from the hurdle package in R, as it specifies one process for zero counts and one process for positive counts, and is commonly used for positively skewed data with lots of zeros (Gonzales-Barron et al. 2010; Hofstetter et al. 2016).

## Results

### Growth Rate

*P. stutzeri* adapted to changes in salinity after ∼330 generations of serial transfer. Both evolution treatments (constant vs. increasing salinity) on average grew faster than the ancestor in the high salt environment. However, populations evolved in the increasing salinity environment grew faster in both the low and high salt environment (Fig. 2A; Salinity p < 0.001; Treatment*Salinity p = 0.052). In addition, populations exposed to the gradual increase in salt, exhibited higher growth rates than those adapted to the constant salt concentration but only when tested at the lower salinity (Fig. 2A; p= 0.0468). Interestingly, both of the evolved populations had larger population sizes in the high salt environment – relative to the low salt environment and to the ancestor (Fig. 2B; p <0.001). This was surprising given the evolved populations grew significantly slower in that environment compared to the low salinity one.

**Fig. 2.**
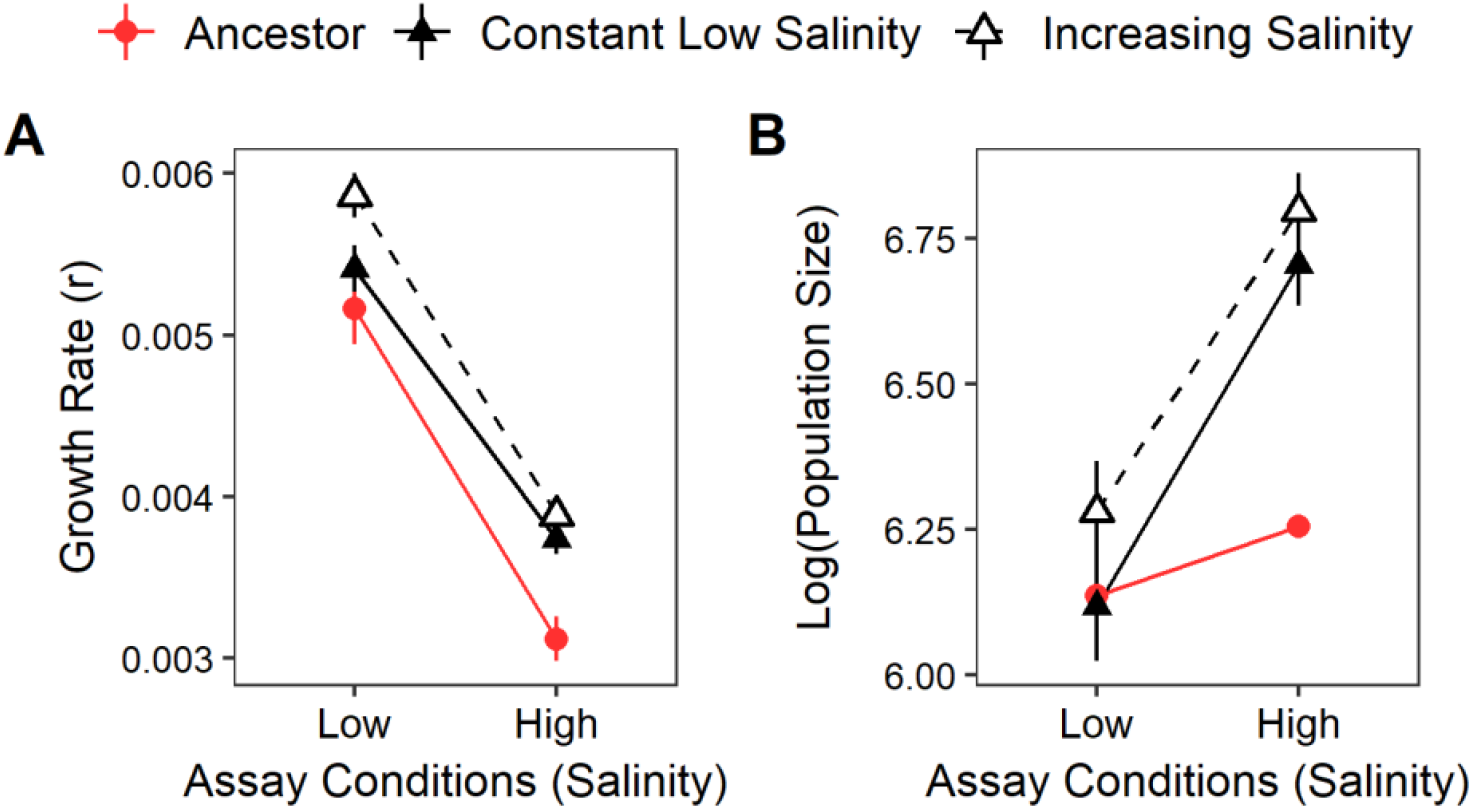
Effects of adaptive evolution. (A) Growth rate and (B) log transformed population size for the ancestor (red circles), constant low salinity (black closed triangles), and increasing salinity evolution treatments (black open circles) in low (1.5%) and high (3%) salinity. The points show the average across the ancestral (n=8) and evolved populations, and the error bars indicate the standard error (constant low, n = 86; increasing n = 94).

### Loss of Transformability

Evolved populations exhibited a significant loss of transformation capacity relative to the ancestor (Fig. 3; constant low salinity, p=0.005; increasing salinity, p=0.02). At the end of the experiment between 39% and 64% of evolved populations – depending on the treatment and test conditions still underwent transformation. In the remaining populations there were no transformants at a detectable level. In addition, there was a striking similarity between the two evolution treatments in terms of how many populations underwent transformation (Fig. 3). In both evolution treatments, there were significantly more populations undergoing transformation when tested at the higher salinity, despite no known difference in genotype (Fig. 3; p <0.001 for both treatments).

**Fig. 3.**
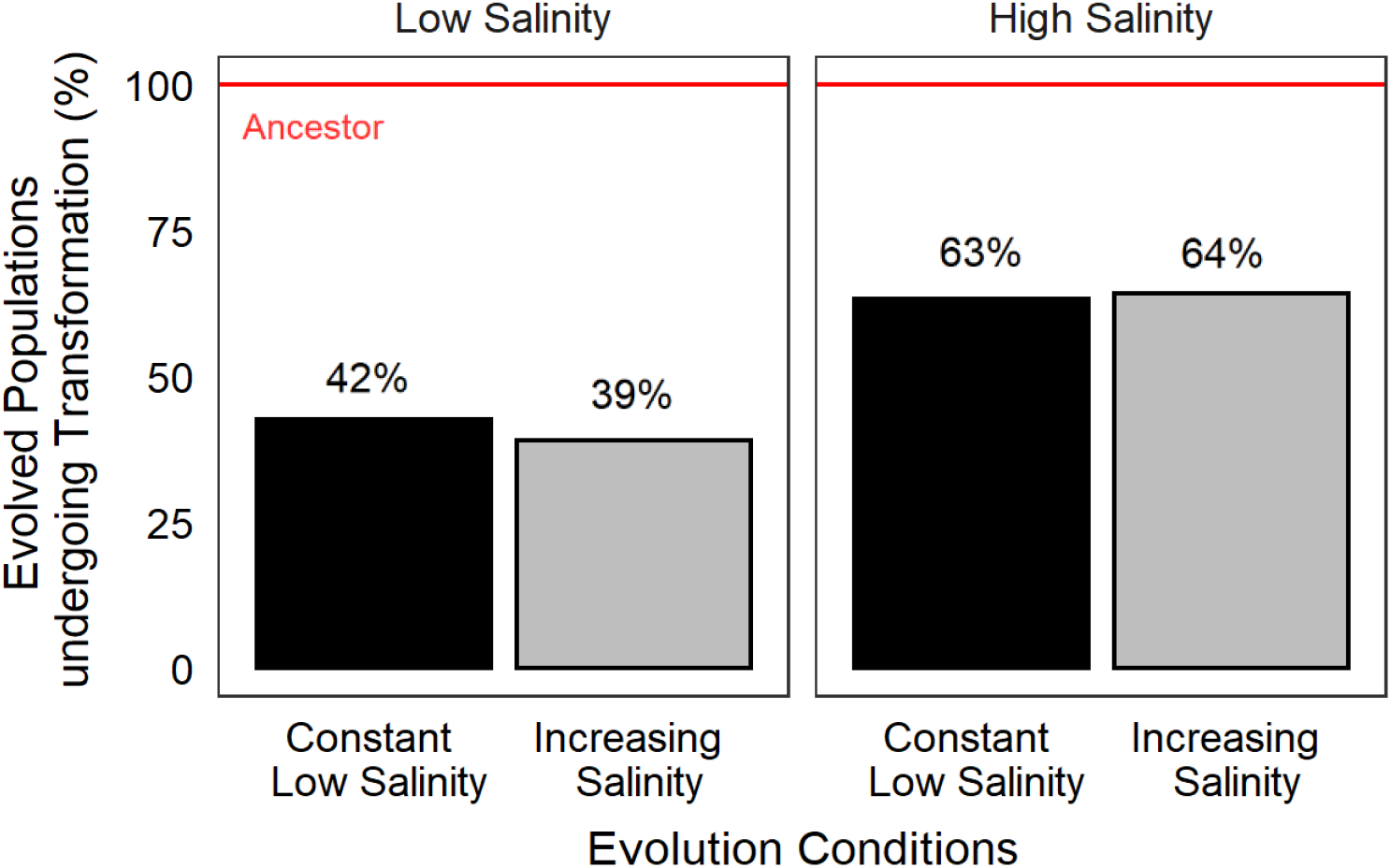
The number of evolved populations with a detectable number of transformants in the low and high salinity test environment for populations evolved in constant or increasing salinities (count data). The red line corresponds to the beginning of the experiment when transformants could be detected in 100% of the ancestral populations.

### Transformants and total cells

There was no relationship between the number of transformants and the number of recipient cells in independently evolving populations (Fig. 4). This trend was true across the treatments, as well as agreeing with preliminary work showing that transformation is not limited by population size in larger populations (Fig. S1A). Therefore, we report the number of transformants standardized by the amount of eDNA (transformation efficiency). However, we also report (in Fig. 5B) the number of transformants standardized by the number of recipient cells (transformation frequency). This is done to account for the fact that, on average, there were significantly more transformants and recipient cells at higher salinities, suggesting the increase in the average number of transformants could be correlated with the increase in the average number of recipient cells – despite there being no evidence of such a correlation within the individual populations.

**Fig. 4.**
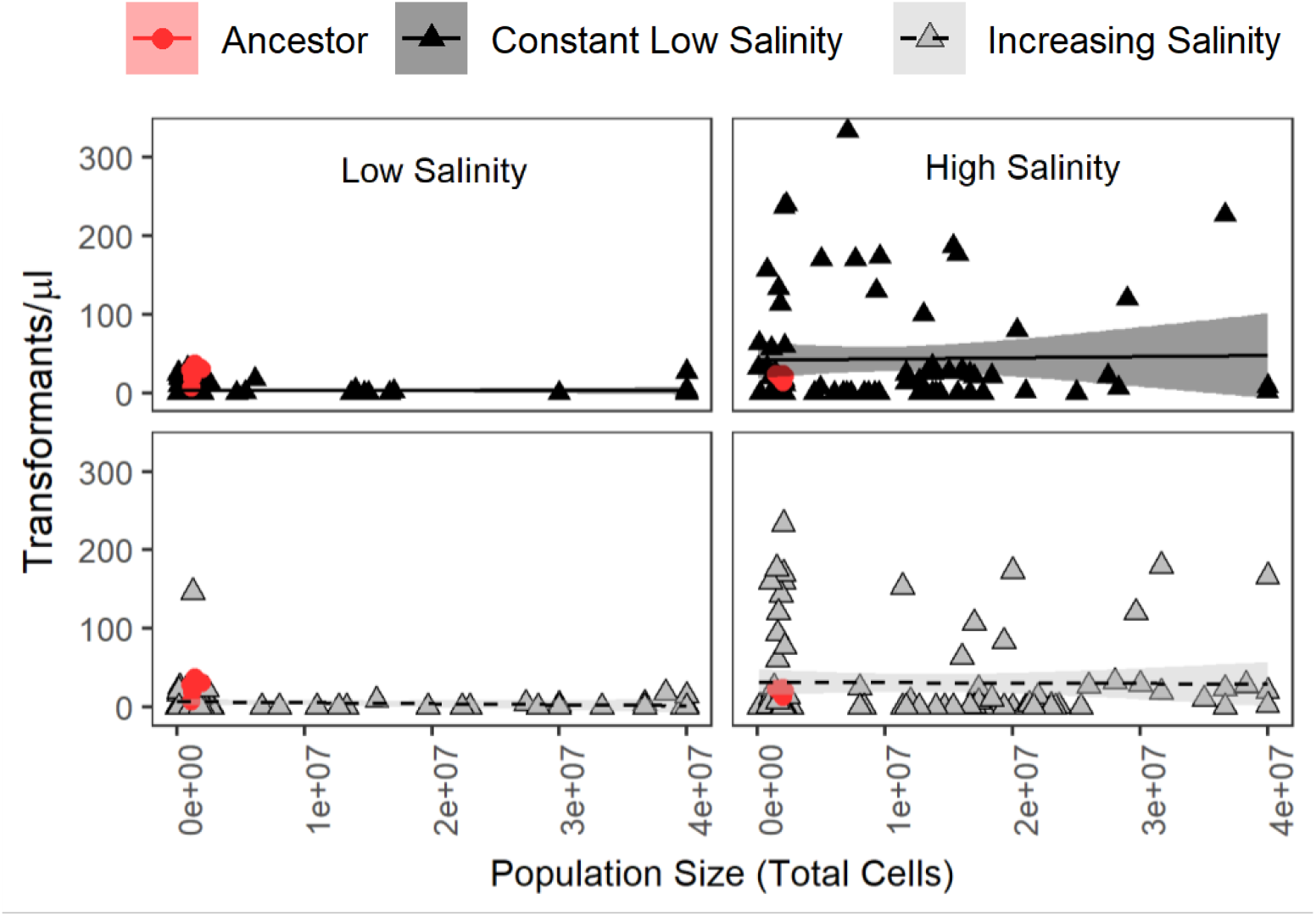
The relationship between transformants and total cells. In the ancestral and evolved populations, the number of transformants does not increase as the population size or total number of cells increases. Each point represents an individual population-level measurement. The linear relationship between transformants and total cells is indicated by the lines with the shaded areas showing the 95% confidence interval (n=8 ancestor, n=86 constant low and n=94 in the increasing salinity treatment).

**Fig. 5.**
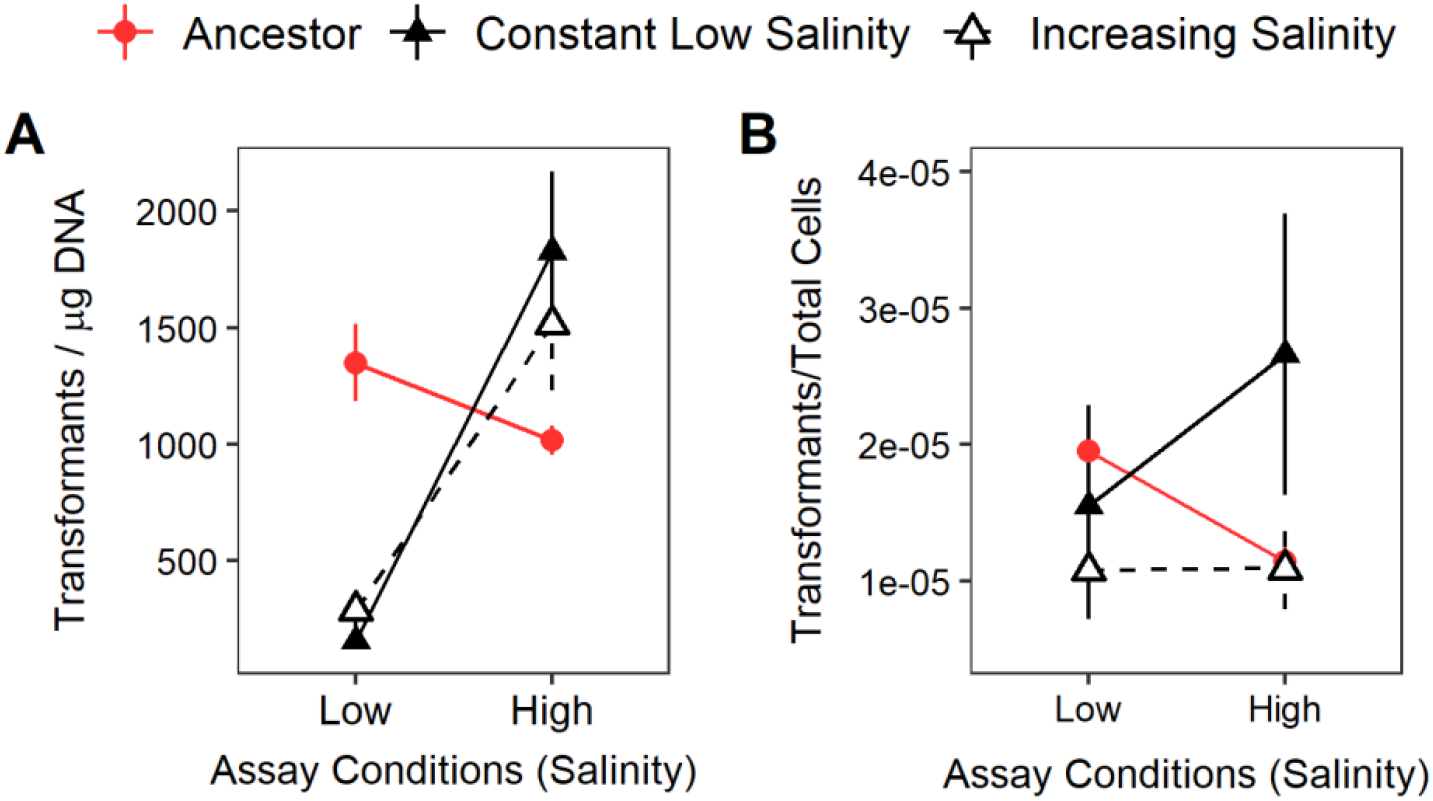
Changes in transformation in response to experimental evolution. (A) Transformation efficiency (transformants/μg DNA) and (B) transformation frequency (transformants/total Cells) for the ancestor (red circles) and evolved populations (constant low salinity = solid black lines; increasing salinity = dashed lines). The points show the average across the ancestral (n=8) and evolved (constant low = 86, increasing =94) populations, and the error bars indicate the standard error (averages include transforming and non-transforming populations).

### High variation in transformation efficiency

At the end of the experiment the transformation efficiency (transformants/μg DNA) was significantly lower in the low salt environment, regardless of the evolution conditions (Fig. 5A; p < 0.0001). However, the populations that transformed eDNA and evolved at constant salinity, did so at a higher efficiency than the ancestor – but only when tested in the high salt environment (Fig. S2; p = 0.0216). When the number of transformants was standardized by the number of recipient cells, there were no statistically significant differences in transformation frequency (Fig. 5B). Although numerically the transformation frequency was highest for populations evolved in constant low salinities and moved to the high salinity environment for the transformation assay.

Evolving with cells lysates or populations of dead cells did not affect transformation efficiency in a uniform manner (Figure 6; Figure S3; see Table S3 for expanded results). In general, the transformation efficiency was higher in the high salt environment. However, standardizing by the population size indicated there was no difference in transformation frequency between the low and high salt environment – as the number of transformants and the number of total cells was larger in the high salt environment (Fig. S4). Moreover, there were no consistent changes in growth rate or population size with the addition of different cell lysates (Fig. S5; see Table S4 and S5 for expanded results). There was a high level of congruency between the two evolution treatments in terms of which populations underwent transformation (Fig. 6; comparing the top and bottom panels). These effects may have appeared early in the experiment, since the two treatments diverged from a single set of evolving populations on day 50 of the experiment.

**Fig. 6.**
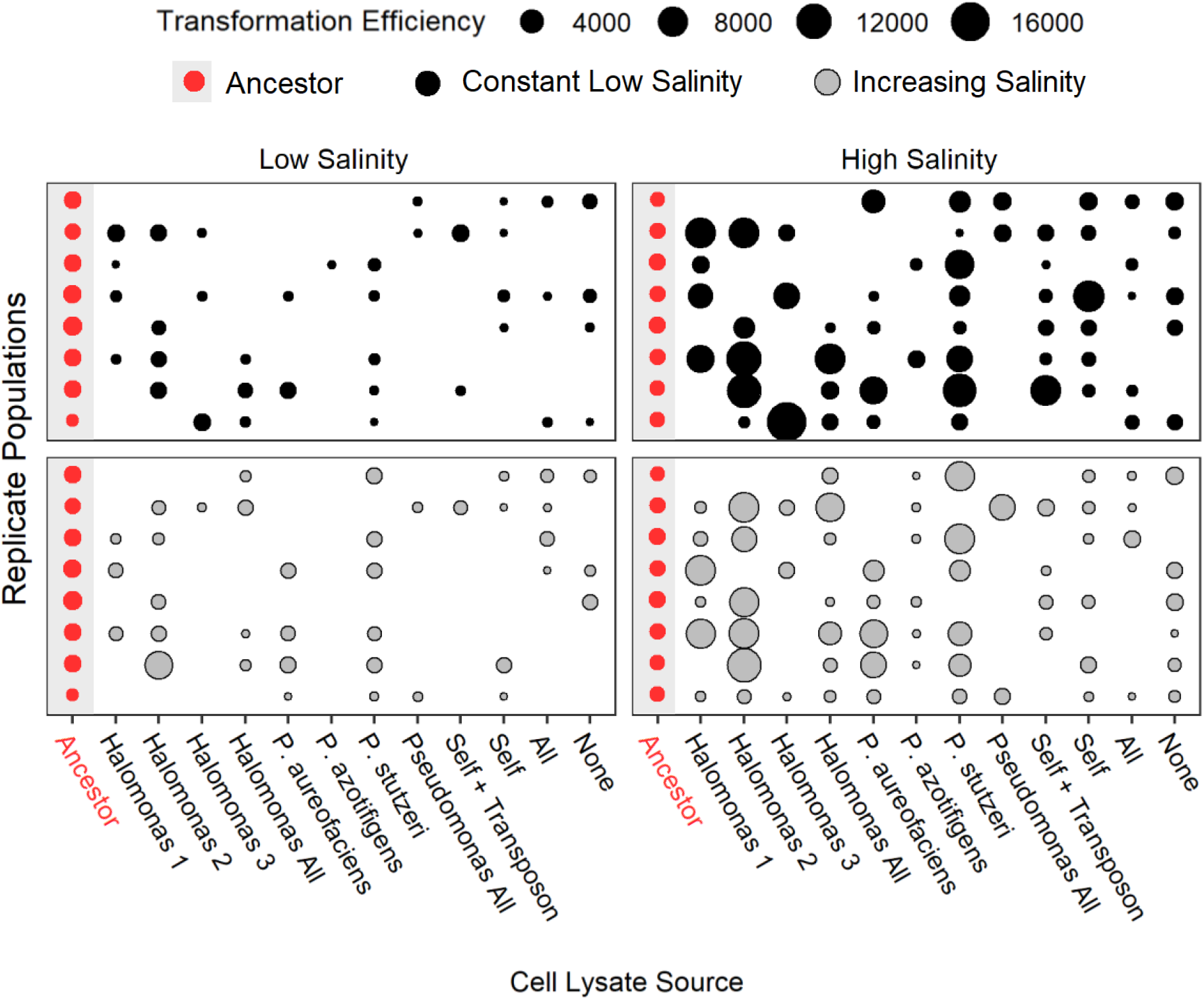
Variation in transformation efficiency. Transformation efficiency for the ancestral and evolved populations in low and high salinities. Higher transformation efficiencies are denoted by larger circles with the populations evolved at a constant low salinity in the top panel and those evolved in increasing salinities in the bottom panel. The cell lysates are shown on the x-axis and the replicate populations on the y-axis corresponding to a 96-well plate layout (n=8 replicates per cell lysate source).

### Trade-off between growth rate and transformation

There was no trade-off between the transformation efficiency and the growth rate (Fig. 7; the same being true for transformation frequency – data not shown). While there may have been tradeoff between growth rate and transformation within individual strains, we were unable to detect such a tradeoff in the population-level measurements conducted here.

**Fig. 7.**
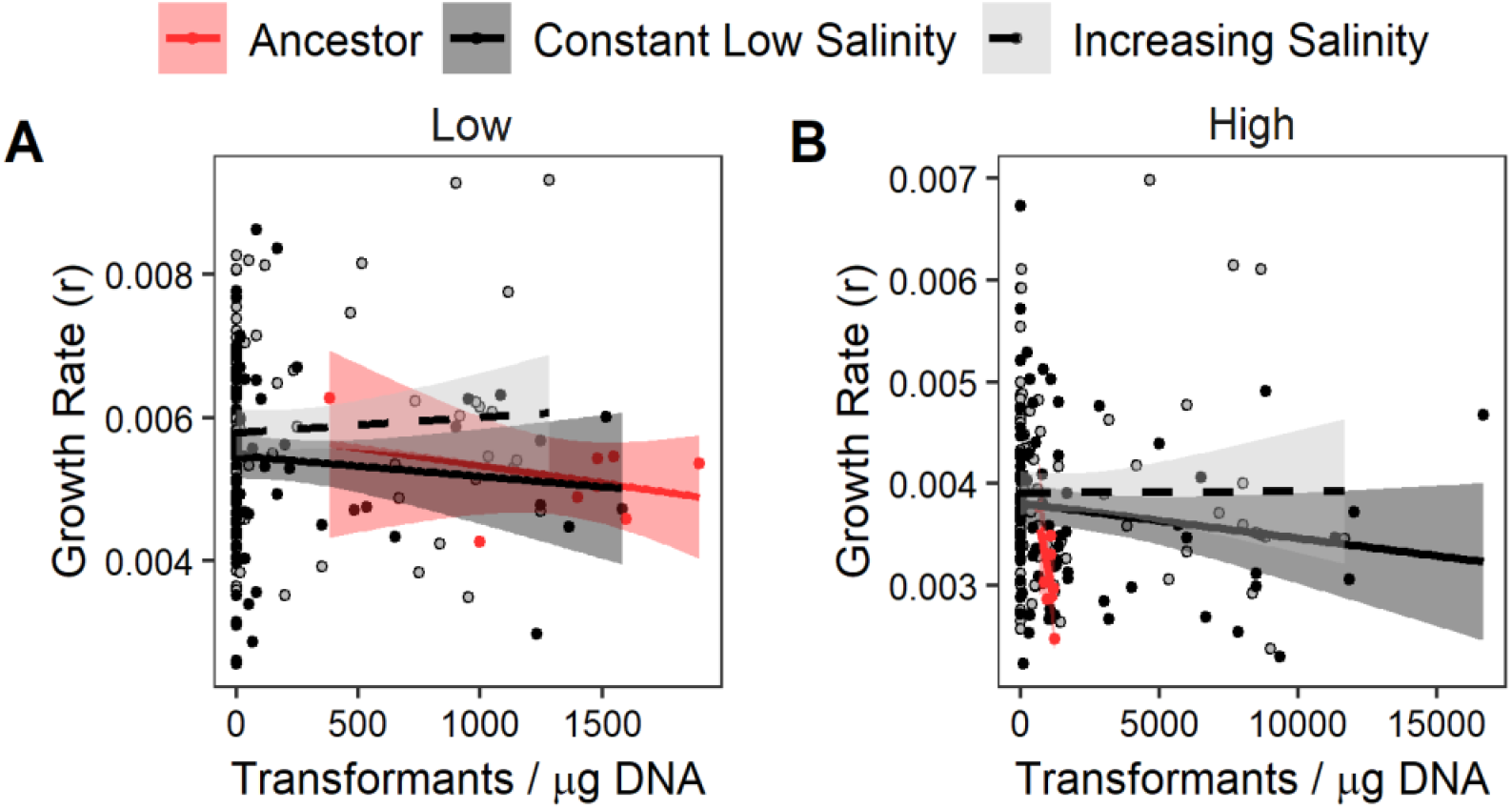
Trade-off between growth rate and transformation efficiency. Average population-level growth rate regressed against the transformation efficiency in the (A) low salinity and (B) high salinity environments. Each point represents an individual population-level measurement. The linear relationship between transformants and total cells is indicated by the line, with the shaded area showing the 95% confidence interval (n=8 ancestor, n=86 constant low and n=94 in the increasing salinity treatment).

## Discussion

Understanding the evolutionary origins and fitness consequences of transformation can shed light on the larger question of why organisms undergo genetic recombination, and to what extent the traits governing selection are themselves selected upon. Here, I conclude that evolving *P. stutzeri* with different sources of eDNA or evolving them in constant versus increasing salinities did not have a large effect on growth rate or transformation efficiency. However, I did find that the transformation capacity – the ability for evolved populations to transform eDNA – changed dramatically over just ∼330 generations or 100 days of serial transfer.

By the end of the experiment, around 50% of the evolved populations did not transform any of the provided extracellular antibiotic resistance genes, although the exact percentage varied from 36-61% depending on the treatment. This was true, regardless of whether the number of transformants was reported as the transformation efficiency (transformants/μg eDNA) or the transformation frequency (transformants/total cells). We report both metrics here to account for differences in average population size in the low versus high salinity environment, even though the total population size only limits transformation in very small populations of *P. stutzeri* (smaller than those reported here; Fig. S1B). Overall, we focus the discussion on the variation in transformation capacity across the evolved lineages, as this is true irrespective of how the data is analyzed (transformation efficiency vs. transformation frequency).

Several other bacterial species, in addition to *P. stutzeri*, undergo transformation irrespective of population density. For instance, *Vibrio parahaemolyticus* and *V. campbellii* both undergo transformation in the absence of quorum sensing which is the ability to regulate gene expression with population size. Meanwhile, their close relative *V. cholerae*, and *Streptococcus* species both require quorum sensing for successful transformation (Shanker and Federle 2017; Simpson et al. 2019). Interestingly, the genetic features that underpin the differences in quorum sensing across *Vibrio* species have yet to be identified. Similarly, different isolates of the same bacterial species often exhibit large differences in their transformation capacity, with the genetic variation underpinning these differences often impossible to discern. For instance, isolates of *P. stutzeri* collected from different soil environments had highly variable transformation frequencies, with about one-third of isolates considered non-transformable (Sikorski et al. 2002). Similar observations have been made in *Vibrio* species that inhabit different environments, and spurred the recent suggestion that transformation may be lost in environments where it is no longer beneficial (Jaskolska et al. 2018; Simpson et al. 2019).

Hence, it is possible that transformation was not maintained in several of the evolved lineages because it did not provide a fitness benefit during experimental evolution. Weak selection for transformation could have been due to osmotic stress, the application of only a mild stress, or due to infrequent fluctuations in the environment. For instance, previous studies found transformation was beneficial when the environment shifted every 2-3 transfers (Perron et al. 2012; Engelmoer et al. 2013), as opposed to every 20-30 transfers as was done in this study. Therefore, an interesting follow-up study would be to compare the distribution of transformation phenotypes after evolving *P. stutzeri* in a constant optimal environment, a constant but very stressful environment, and a rapidly fluctuating environment – to better understand how stress, or fluctuations in stress, shape the evolution of transformation.

Another possibility is that transformation only provides a fitness benefit in response to very specific stressors. For instance, several studies have found that transformation was beneficial when evolving populations where exposed to periodic inputs of sub-inhibitory concentrations of antibiotics (Perron et al. 2012; Engelmoer et al. 2013). Because transformation allows the reversible integration of resistance genes, and antibiotics are usually transiently present in the environment, transformation could be a mechanism well-suited to handling antibiotic stress. In contrast, transformation may be less beneficial in response to stressors like changes in osmotic pressure which are encoded by large and connected gene networks. In general, more work needs to be done on the specific stressors that transformation confers a benefit to, as prokaryotic genes appear to adapt to either vertical or horizontal transmission, meaning that not all processes may be well-adapted to evolve via horizontal gene transfer (Novick and Doolittle 2020).

A final consideration is the role of osmotic pressure in altering the efficiency of eDNA uptake. Populations adapting to high osmolarity environments generally have a large fraction of mutations in genes associated with cell wall synthesis (Cesar et al. 2020). Therefore, it could be that changes in the cell wall altered the pilus structure which captures eDNA from the environment (Graupner et al. 2000). *P. stutzeri* has two pili that interact to regulate transformation. The type IV pilus acquires eDNA from the environment, while the second pilus is believed to translocate eDNA into the cytoplasm and when knocked out decreases transformation ∼90% (Graupner et al. 2001). Therefore, changes in the cell wall in response to salt stress could have altered the interaction between these two pili, creating the gradient of transformation capacity evident in the evolved lineages. Follow-up investigations which involve whole genome sequencing, will hopefully elucidate if mutations in osmoregulatory genes could have altered transformation capacity.

Despite evidence of high variation in transformation capacity in many bacterial lineages, very few studies have quantified changes in transformation after experimental evolution. To date, six experimental evolution studies have focused on the fitness effects of transformation but only one study has quantified transformation before and after experimental evolution. In that one study, transformation did not provide a fitness benefit and the evolved lineages repeatedly lost the capacity to undergo transformation (Bacher et al. 2006). Several other studies have identified transformation-mediated fitness benefits (primarily in stressful environments), but none of them quantified the prevalence of transformation at the end of the experiment (Table 1).

Moving forward, quantifying transformation after adaptation could help disentangle the benefits of transformation from the overall benefits of competence – which is the physiological state a bacterium must enter to undergo transformation and is often part of a larger stress response. For instance, Pseudomonads also use the Type IV pilus for flagellum-independent movement or twitching motility (Graupner et al. 2001). While *Bacillus subtilis*, a well-studied soil-dweller, upregulates transformation as part of a general stress response prompted by DNA damage or antibiotics (Claverys et al. 2006). Therefore, future studies that quantify changes in transformation efficiency during and after experimental evolution, will be critical in disentangling the specific benefits of transformation within the larger regulatory network of competence.

Overall, this study provides novel experimental evidence that the ability to undergo transformation can change over relatively short timescales and may be more plastic across space and time than is generally accepted. The substantial decrease in transformation efficiency in the low salt environment where *P. stutzeri* evolved, suggests that transformation did not provide a fitness benefit during salt adaptation. Taken together, this work suggests that transformation is an adaptive, selectable trait, that may increase or decrease rapidly in response to selection.

---

**Fig. S1.**
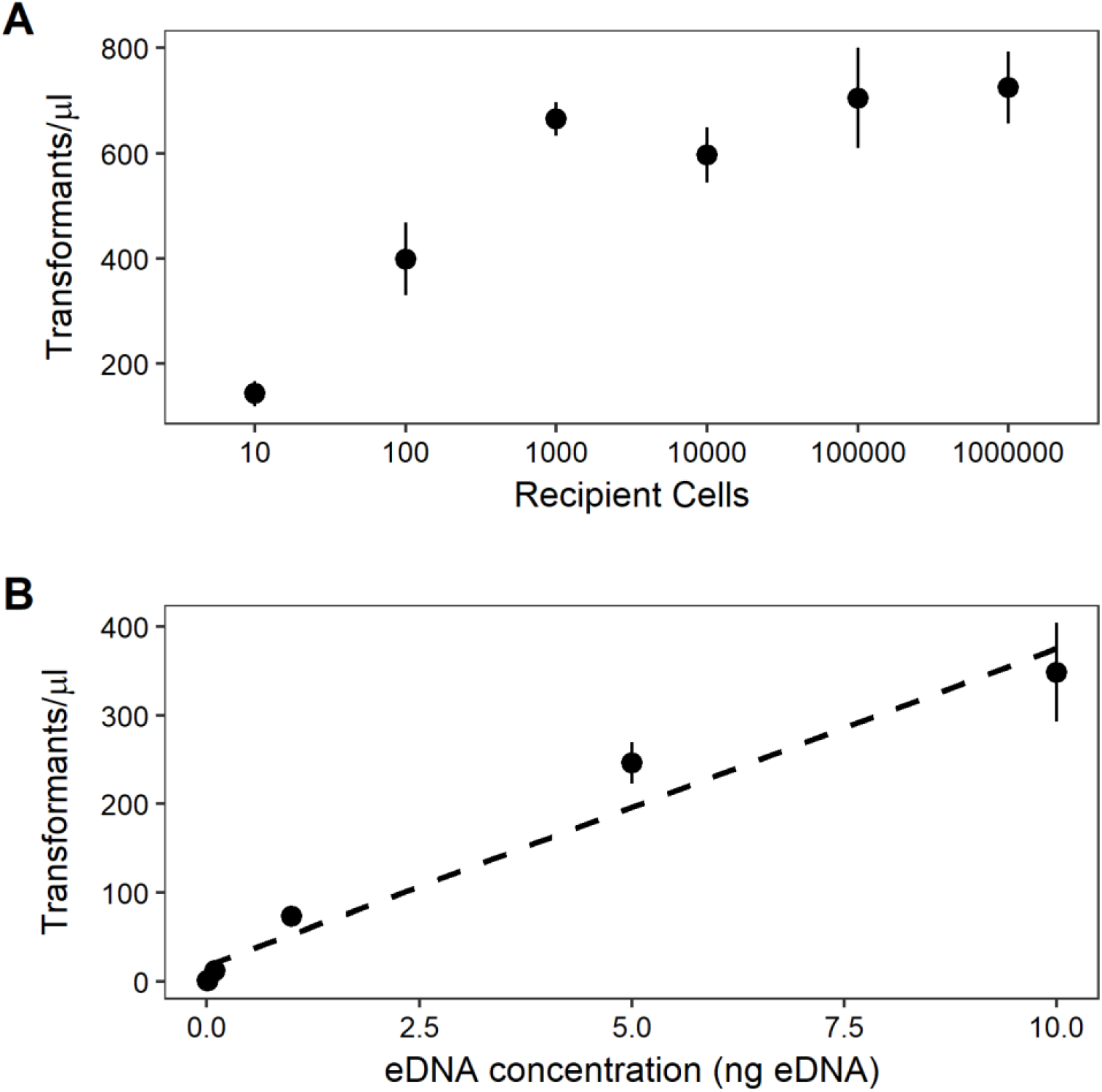
Preliminary transformation assays. (A) The effect of total population size on the number of transformants when the eDNA concentration is held constant. (B) The relationship between the concentration of eDNA and the number of transformants under laboratory conditions (0.001, 0.01, 0.1, 1, 5, 10 ng/μl eDNA) – in large populations (∼1,000,000 recipient cells).

**Fig. S2.**
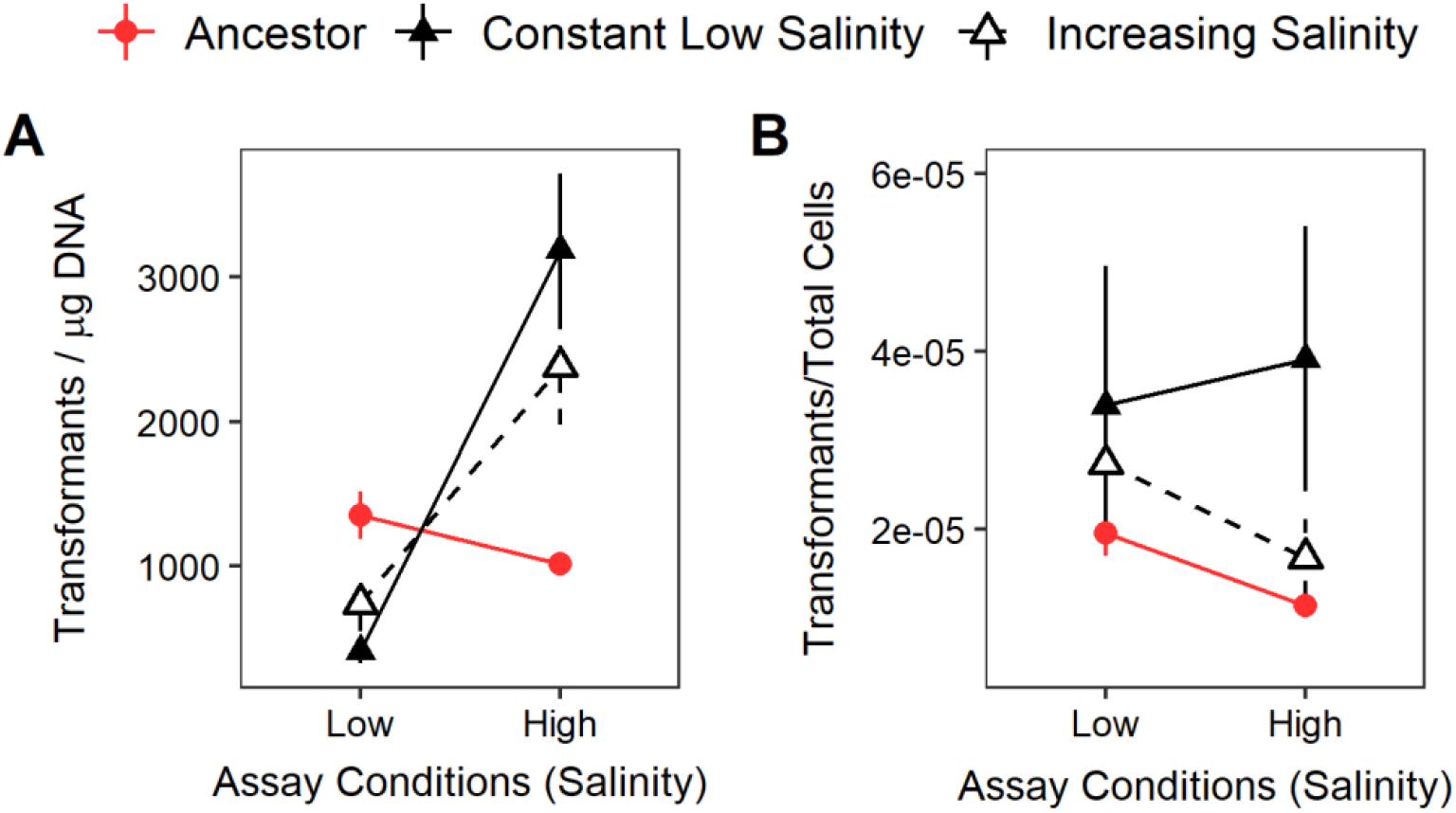
Transformability in the populations that transformed eDNA. (A) Transformation efficiency (transformants/μg DNA) and (B) transformation frequency (transformants/total Cells) for the ancestor (red circles) and evolved populations (constant low salinity = solid black lines; increasing salinity = dashed lines). The points show the average across the ancestral (n=8) and evolved (constant low = 86, increasing =94) populations, and the error bars indicate the standard error (averages include only populations that underwent transformation).

**Fig. S3.**
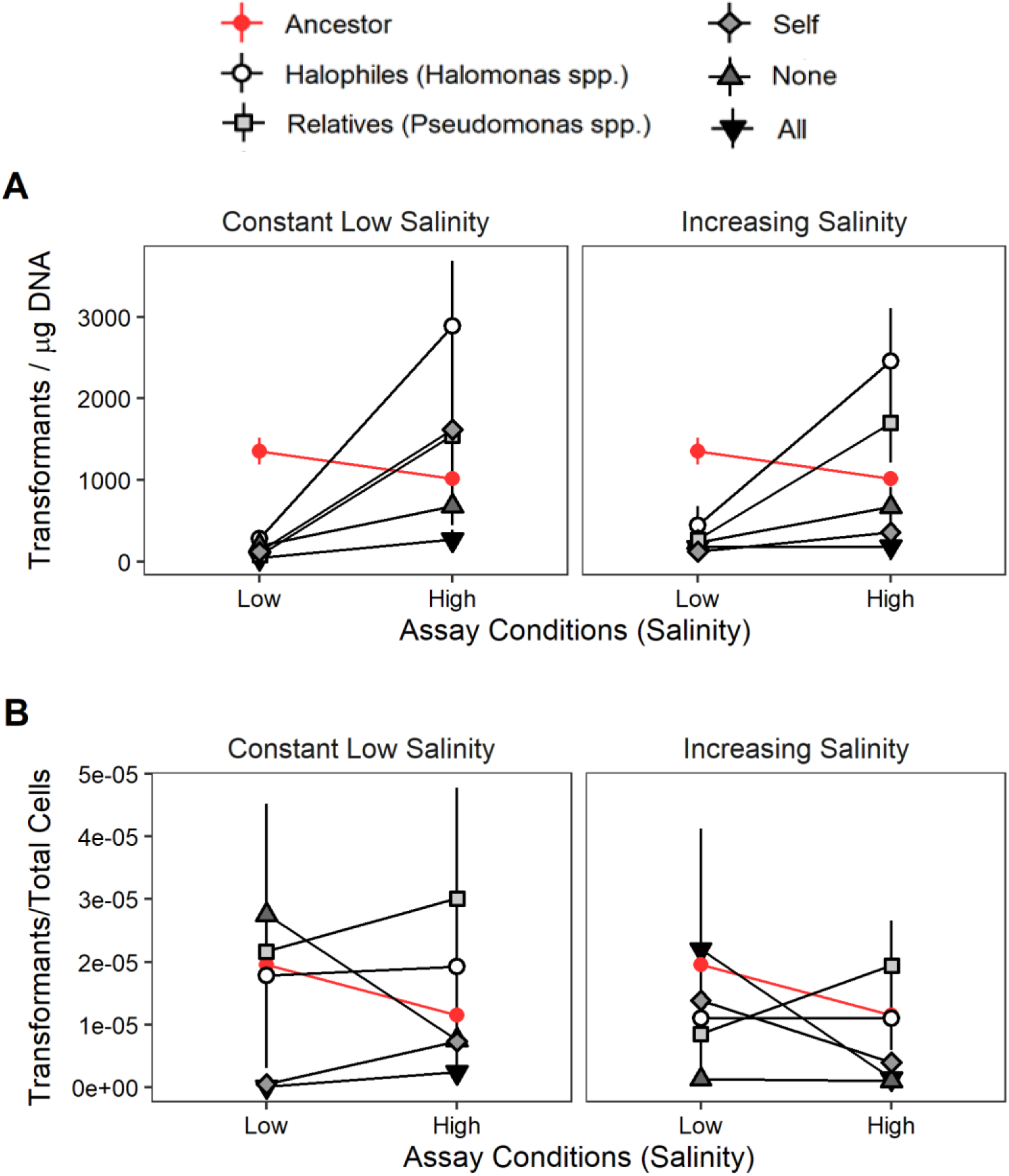
Effect of cell lysates on transformation. (A) Transformation efficiency, and (B) transformation frequency for each cell lysate treatment. The ancestor (red circles), the constant low salinity (left panel) and the increasing salinity treatments (right panel, dashed lines) are shown at low and high salinity (n=8 replicate populations across treatments, excluding treatments that went extinct, see table S4.2).

**Fig. S4.**
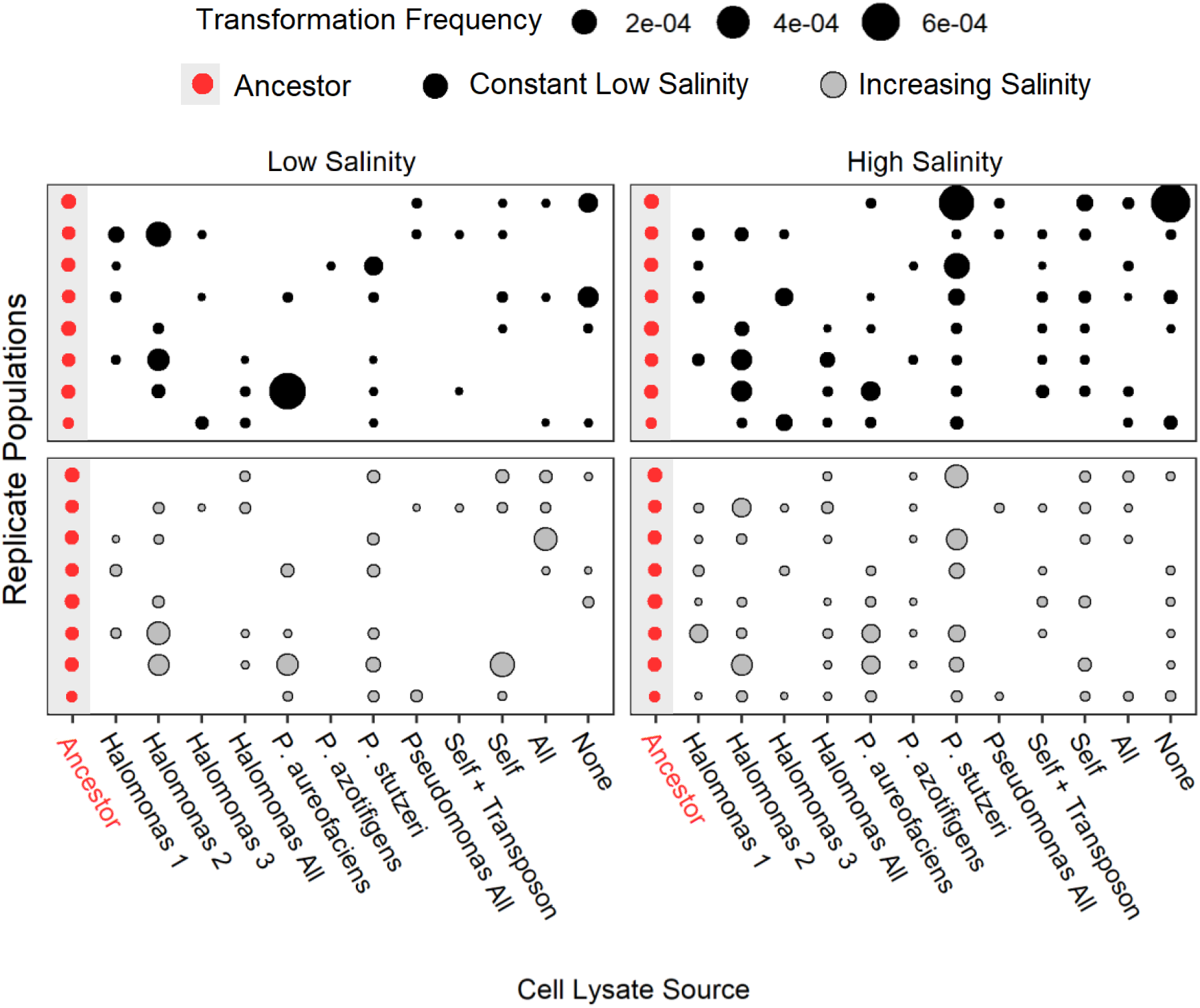
Variation in transformation frequency. Transformation frequency for the ancestral and evolved populations in low and high salinities. Higher transformation frequencies are denoted by larger circles with the populations evolved at a constant low salinity in the top panel and those evolved in increasing salinities in the bottom two panels. The cell lysates are shown on the x-axis and the replicate populations on the y-axis corresponding to a 96-well plate (n=8 replicates per cell lysate).

**Fig. S5:**
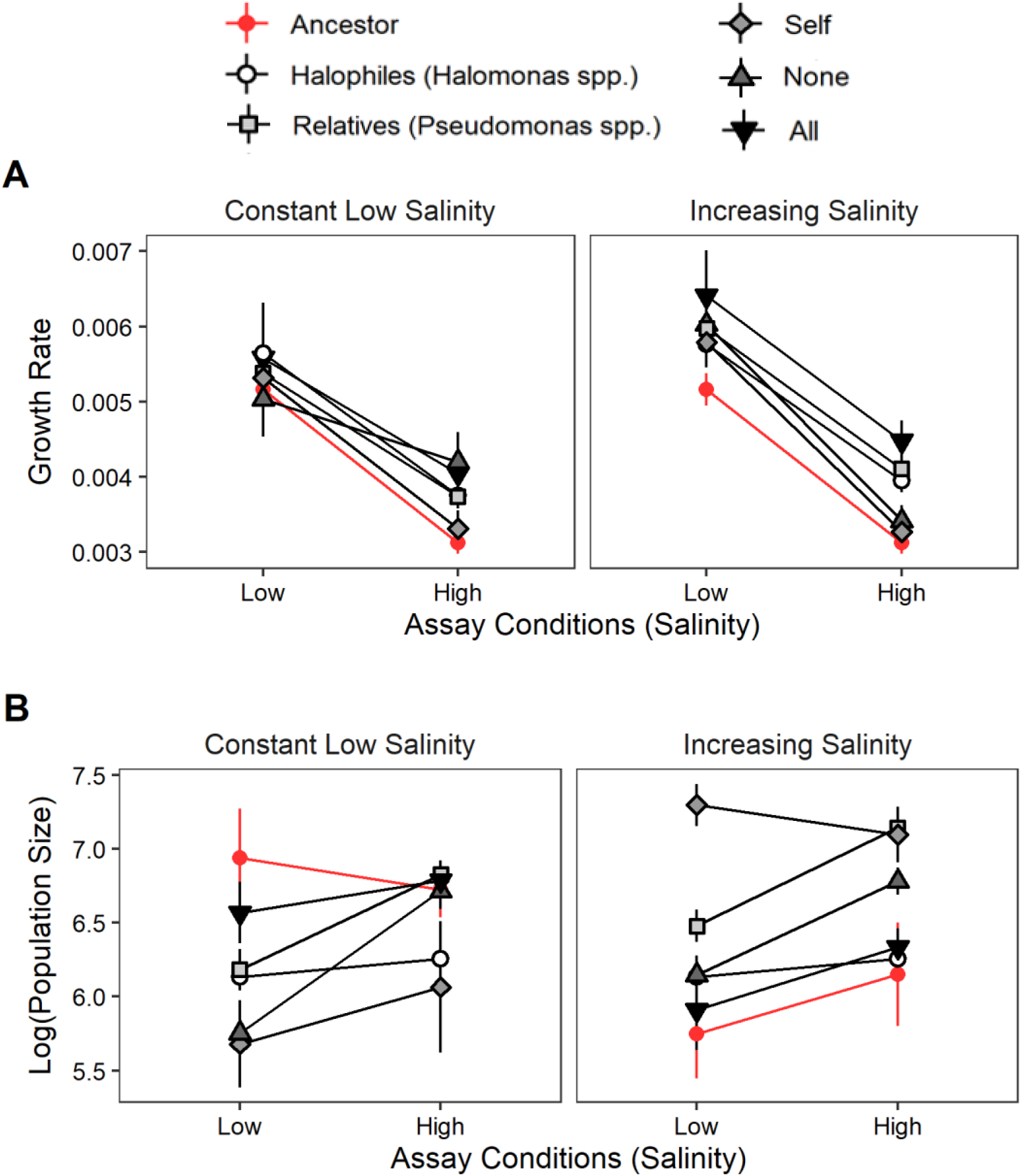
Effect of cell lysates on growth rate and population size. (A) Growth rate, and (B) log transformed population size for each cell lysate treatment. The ancestor (red circles), the constant low salinity (left panel) and the increasing salinity treatments (right panel, dashed lines) are shown at low and high salinity (n=8 replicate population across treatments).

**Table S1.** Summary of Cell lysate sources. The halophiles were collected from salt springs in the Namib dessert. The conductivity of the spring at the time of collection is listed for *Halomonas spp*. The *Pseudomonas* relatives were purchased from the German culture collection (accessions number listed).

| Strain | Cell Lysate Category | Strain Source | Conductivity (mS/cm) |
| --- | --- | --- | --- |
| Halomonas spp. 1 (*Halomonas taeanensis*) | Halophile | Namib Springs Aub Canyon | 7.3 |
| Halomonas spp. 2 (*Halomonas*) | Halophile | Namib Springs Kai-As | 1 |
| Halomonas spp. 3 (*Halomonas*) | Halophile | Namib Springs Swartmodder | 190 |
| Halomonas spp. All (1+2+3) | Halophile |  |  |
| *Pseudomonas chlororaphis* subsp. *aureofaciens* | Relative | <u>DSM6698</u> |  |
| *Pseudomonas azotifigens* | Relative | <u>DSM17556</u> |  |
| *Pseudomonas stutzeri* JM300 | Relative | <u>DSM10701</u> |  |
| *Pseudomonas* spp. All | Relative |  |  |
| Self-Lysate + Transposon (*P. stutzeri* + miniTn7) | Self | Baltrus lab |  |
| Self-Lysate (*P. stutzeri*) | Self | Baltrus lab |  |
| None | None |  |  |
| All | All | A combination of all strains listed above |  |

**Table S2.**
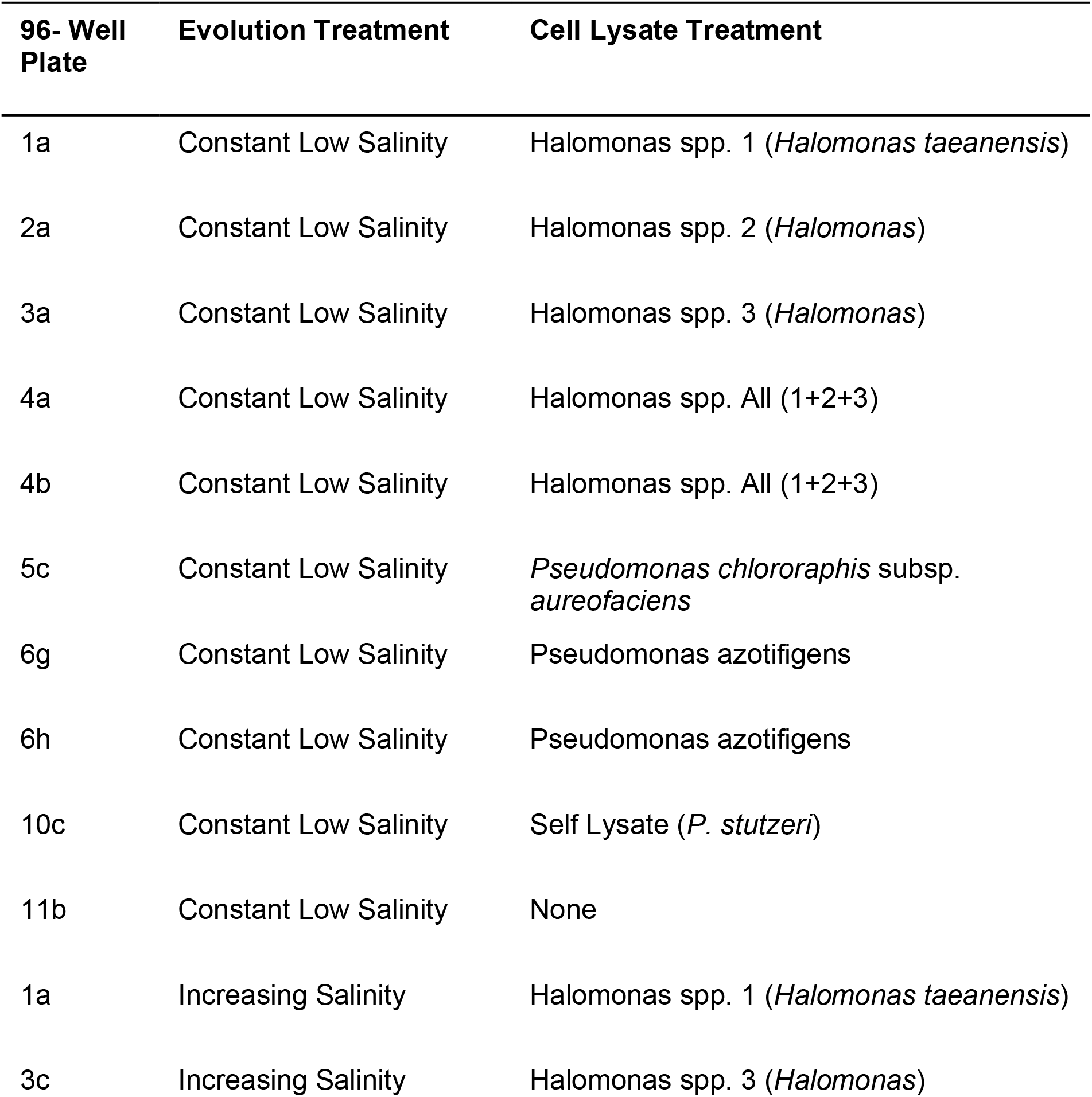
Populations that did not revive. List of populations that did not revive at the end of the experiment. One population was contaminated (3c).

**Table S3.** Expanded transformation efficiency results. Letters for pairwise comparisons across the cell lysate treatments with p-values listed at the bottom of the column.

|  | Transformation Efficiency |  |  |  |
| --- | --- | --- | --- | --- |
|  | Constant Salinity |  | Increasing Salinity |  |
|  | 1.50% | 3.00% | 1.50% | 3.00% |
| Ancestor | a | a | b | a |
| Self Lysate | b | a | a | a |
| No Lysate | b | a | ab | a |
| Halophile Lysate | b | a | ab | a |
| Relatives Lysate | b | a | a | a |
| All spp. | b | a | ab | a |
|  | p<0.0001 | p=0.055 | p=0.0207 | p=0.0506 |

**Table S4.** Expanded growth rate results. Letters for pairwise comparisons across the cell lysate treatments with p-values listed at the bottom of the column.

|  | Growth Rate (r) |  |  |  |
| --- | --- | --- | --- | --- |
|  | Constant Salinity |  | Increasing Salinity |  |
|  | 1.50% | 3.00% | 1.50% | 3.00% |
| Ancestor | a | a | a | a |
| Self Lysate | a | ab | ab | ab |
| No Lysate | a | b | a | abc |
| Halophile Lysate | a | b | ab | bc |
| Relatives Lysate | a | b | b | c |
| All spp. | a | b | ab | c |
|  | p=0.697 | p<0.001 | p=0.08 | p<0.001 |

**Table S5.**
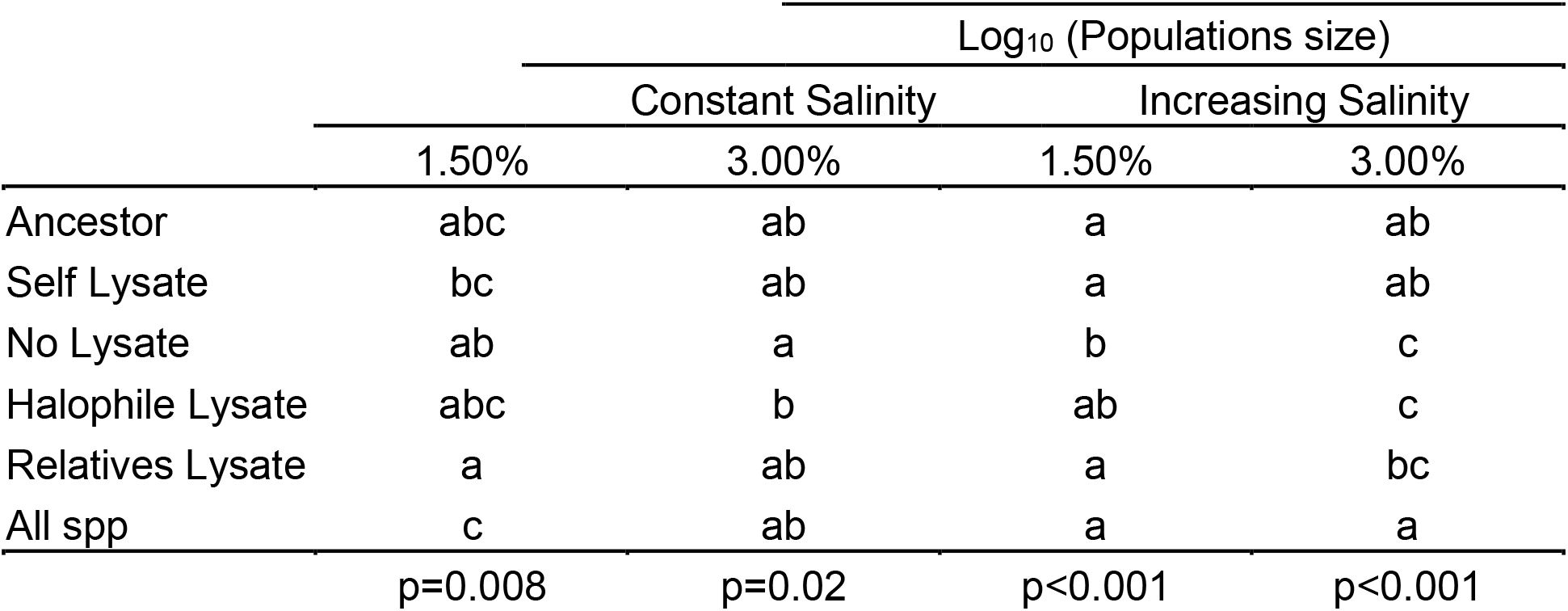
Expanded population size results. Letters for pairwise comparisons across the cell lysate treatments with p-values listed at the bottom of the column.

## References

Ambur OH, Engelstädter J, Johnsen PJ, et al (2016) Steady at the wheel: Conservative sex and the benefits of bacterial transformation. Philos Trans R Soc B Biol Sci 371:. doi: 10.1098/rstb.2015.0528

Bacher JM, Metzgar D, Valerie de C-L (2006) Rapid Evolution of Diminished Transformability in Acinetobacter baylyi □. J Bacteriol 188:8534–8542. doi: 10.1128/JB.00846-06

Baltrus DA (2013) Exploring the costs of horizontal gene transfer. Trends Ecol Evol 28:489–495. doi: http://dx.doi.org/10.1016/j.tree.2013.04.002

Baltrus DA, Guillemin K, Phillips PC (2007) Natural Transformation increases the rate of adaptation in the human pathogen Helicobacter pylori. Evolution (N Y) 62:39–49. doi: 10.1111/j.1558-5646.2007.00271.x

Barton NH, Charlesworth B (1998) Why Sex and Recombination ? 281:1986–1991

Carvalho G, Fouchet D, Danesh G, et al (2019) Bacterial transformation buffers environmental fluctuations through the reversible integration of mobile genetic elements. 1–12

Cesar S, Anjur-dietrich M, Yu B, et al (2020) Bacterial Evolution in High-Osmilarity Environments. 11:1–17

Charpentier X, Polard P, Claverys J (2012) Induction of competence for genetic transformation by antibiotics : convergent evolution of stress responses in distant bacterial species lacking SOS ? Curr Opin Microbiol 15:570–576. doi: 10.1016/j.mib.2012.08.001

Claverys J, Prudhomme M, Martin B (2006) Induction of Competence Regulons as a General Response to Stress in Gram-Positive Bacteria. doi: 10.1146/annurev.micro.60.080805.142139

Cooper TF (2007) Recombination speeds adaptation by reducing competition between beneficial mutations in populations of Escherichia coli. PLoS Biol 5:1899–1905. doi: 10.1371/journal.pbio.0050225

Engelmoer DJP, Donaldson I, Rozen DE (2013) Conservative Sex and the Benefits of Transformation in Streptococcus pneumoniae. PLoS Pathog 9:1–7. doi: 10.1371/journal.ppat.1003758

Gonzales-Barron U, Kerr M, Sheridan JJ, Butler F (2010) Count data distributions and their zero-modified equivalents as a framework for modelling microbial data with a relatively high occurrence of zero counts. Int J Food Microbiol 136:268–277. doi: 10.1016/j.ijfoodmicro.2009.10.016

Graupner S, Frey V, Hashemi R, et al (2000) Type IV pilus genes pilA and pilC of Pseudomonas stutzeri are required for natural genetic transformation, and pilA can be replaced by corresponding genes from nontransformable species. J Bacteriol 182:2184–2190. doi: 10.1128/JB.182.8.2184-2190.2000

Graupner S, Weger N, Sohni M, Wackernagel W (2001) Requirement of Novel Competence Genes pilT and pilU of Pseudomonas stutzeri for Natural Transformation and Suppression of pilT Deficiency by a Hexahistidine Tag on the Type IV Pilus Protein PilAI. 183:4694–4701. doi: 10.1128/JB.183.16.4694

Guiral S, Mitchell TJ, Martin B, Claverys J (2005) Competence-programmed predation of noncompetent cells in the human pathogen Streptococcus pneumoniae : Genetic requirements. PNAS

Hofstetter H, Dusseldorp E, Zeileis A, Schuller AA (2016) Modeling Caries Experience: Advantages of the use of the hurdle model. Caries Res 50:517–526. doi: 10.1159/000448197

Jaskolska M, Stutzmann S, Stoudmann C, Blokesch M (2018) QstR-dependent regulation of natural competence and type VI secretion in Vibrio cholerae. Nucleic Acids Res 46:10619–10634. doi: 10.1093/nar/gky717

Johnston C, Martin B, Fichant G, et al (2014) Bacterial transformation: Distribution, shared mechanisms and divergent control. Nat. Rev. Microbiol. 12

Kim Y, Orr HA (2005) Adaptation in sexuals vs. asexuals: Clonal interference and the Fisher-Muller model. Genetics 171:1377–1386. doi: 10.1534/genetics.105.045252

Koonin E V. (2016) Horizontal gene transfer: essentiality and evolvability in prokaryotes, and roles in evolutionary transitions. F1000Research 5:1805. doi: 10.12688/f1000research.8737.1

Lynch M, rger R, Buthcer D, Gabriel W (1993) The mutation meltdown in asexual populations. JHered 84:339–344

Mcleman A, Sierocinski P, Hesse E, et al (2016) No effect of natural transformation on the evolution of resistance to bacteriophages in the Acinetobacter baylyi model system. Nat Sci Reports 6–11. doi: 10.1038/srep37144

Mell JC, Redfield RJ (2014) Natural Competence and the Evolution of DNA Uptake Specificity COMPETENCE. 196:1471–1483. doi: 10.1128/JB.01293-13

Michod RE, Wojciechowski MF, Hoezler MA (1988) DNA repair and the evolution of transformation in the bacterium Bacillus subtilis. Genetics 118:31–39. doi: 10.1046/j.1420-9101.1994.7020147.x

Moradigaravand D, Engelsta J (2013) The Evolution of Natural Competence : Disentangling Costs and Benefits of Sex in Bacteria. 182:. doi: 10.1086/671909

Novick A, Doolittle WF (2020) Horizontal persistence and the complexity hypothesis. Biol Philos 35:1–22. doi: 10.1007/s10539-019-9727-6

Palmer ND, Cartwright RA (2018) Strong Episodic Selection for Natural Competence for Transformation Due to Host-Pathogen Dynamics. 1–22

Perron GG, Lee AEG, Wang Y, et al (2012) Bacterial recombination promotes the evolution of multi-drug-resistance in functionally diverse populations. Proc R Soc B Biol Sci 279:1477–1484. doi: 10.1098/rspb.2011.1933

Redfield RJ (1993) Genes for breakfast: The have-your-cake and-eat-lt-too of bacterial transformation. J Hered 84:400–404. doi: 10.1093/oxfordjournals.jhered.a111361

Redfield RJ (1988) Evolution of Bacterial Transformation: Is Sex With Dead Cells Ever Better Than No Sex at All?

Redfield RJ, Schragt MR, Dean AM (1997) The Evolution of Bacterial Transformation: Sex With Poor Relations. 38:

Seitz P, Blokesch M (2013) Cues and regulatory pathways involved in natural competence and transformation in pathogenic and environmental Gram-negative bacteria. FEMS Microbiol Rev 37:336–363. doi: 10.1111/j.1574-6976.2012.00353.x

Shanker E, Federle MJ (2017) Quorum sensing regulation of competence and bacteriocins in Streptococcus pneumoniae and mutans. Genes (Basel) 8:. doi: 10.3390/genes8010015

Sikorski J, Teschner N, Wackernagel W (2002) Highly different levels of natural transformation are associated with genomic subgroups within a local population of Pseudomonas stutzeri from soil. Appl Environ Microbiol 68:865–873. doi: 10.1128/AEM.68.2.865-873.2002

Simpson CA, Podicheti R, Rusch DB, et al (2019) Diversity in Natural Transformation Frequencies and Regulation across Vibrio Species. 10:1–16

Sinha S, Mell J, Redfield R (2013) The availability of purine nucleotides regulates natural competence by controlling translation of the competence activator Sxy. 88:1106–1119. doi: 10.1111/mmi.12245

Smith BA, Dougherty KM, Baltrus DA (2014) Complete Genome Sequence of the Highly Transformable Pseudomonas stutzeri Strain 28a24. Genome Announc 2:2014. doi: 10.1128/genomeA.00543-14

Soucy SM, Huang J, Gogarten JP (2015) Horizontal gene transfer: Building the web of life. Nat Rev Genet 16:472–482. doi: 10.1038/nrg3962

Takeuchi N, Kaneko K, Koonin E V. (2014) Horizontal gene transfer can rescue prokaryotes from Muller’s ratchet: Benefit of DNA from dead cells and population subdivision. G3 Genes, Genomes, Genet 4:325–339. doi: 10.1534/g3.113.009845

Veening JW, Blokesch M (2017) Interbacterial predation as a strategy for DNA acquisition in naturally competent bacteria. Nat Rev Microbiol 15:621–629. doi: 10.1038/nrmicro.2017.66

